# Propagation of BOLD activity reveals task-dependent directed interactions across human visual cortex

**DOI:** 10.1101/172452

**Authors:** Nicolás Gravel, Remco J. Renken, Ben M. Harvey, Gustavo Deco, Frans W. Cornelissen, Matthieu Gilson

## Abstract

It has recently been shown that large-scale propagation of blood-oxygen level dependent (BOLD) activity is constrained by anatomical connections and reflects transitions between behavioral states. It remains to be seen, however, if the propagation of BOLD activity can also relate to the brain anatomical structure at a more local scale. Here, we hypothesized that BOLD propagation reflects structured neuronal activity across early visual field maps. To explore this hypothesis, we characterize the propagation of BOLD activity across V1, V2 and V3 using a modeling approach that aims to disentangle the contributions of local activity and directed interactions in shaping BOLD propagation. It does so by estimating the effective connectivity (EC) and the excitability of a noise-diffusion network to reproduce the spatiotemporal covariance structure of the data. We apply our approach to 7T fMRI recordings acquired during resting state (RS) and visual field mapping (VFM). Our results reveal different EC interactions and changes in cortical excitability in RS and VFM, and point to a reconfiguration of feedforward and feedback interactions across the visual system. We conclude that the propagation of BOLD activity has functional relevance, as it reveals directed interactions and changes in cortical excitability in a task-dependent manner.

## 1 Introduction

Neuronal connections among cortical areas can be observed at a variety of scales, from laminar circuits to cortico-thalamic and cortico-cortical connections. Together they form a complex and intricate set of connections that serve as pathways for signal transmission and processing. The anatomy and function of such connections has been a focus of much research during the last decades [5, 83]. Thanks to non-invasive forms of neuronal recordings, like functional magnetic resonance imaging fMRI (see Raichle (2000) [68] for a detailed review), the link between structural and functional connectivity in the human brain has begun to be unravelled in-vivo. However, due to the multiple physiological mechanisms contributing to the BOLD signal (i.e. metabolic, vascular, neuronal, etc), its limited temporal resolution, and high noise level in its measurements [82, 11], it is still difficult to quantify and interpret this relationship link using fMRI. Several strategies have been proposed to address these issues and infer the efficacy with which anatomical connections modulate interactions between brain regions — referred to as effective connectivity (EC) [17, 22, 76, 21]. However, most such strategies rely on the assumption of temporal precedence ‒the temporal resolution of the BOLD signal and its measurement must be sufficient to capture the time scale of modulatory influences. Crucially, the observed responses should reflect temporal dependencies within the system under scrutiny, which may not be the case if hemodynamic response delays differ between regions [2, 4, 29, 26].

Nevertheless, for whole-brain fMRI recordings, it has been recently shown that the propagation of BOLD activity reflects the transition between different behavioral states and may partly map to anatomical paths [58, 57]. This suggests a meaningful relationship between BOLD activity propagation and the modulation of communication between distant brain regions. Therefore, we ask if the propagation of BOLD activity can also reveal relevant aspects of brain activity at a more local scale, such as that of early cortical visual field maps, which are richly interconnected and where regional variation in the hemodynamic response is less pronounced [26, 48]. In the present study, we hypothesize that the propagation of BOLD activity across early cortical visual field maps V1, V2 and V3 reflects structured neuronal activity (i.e. modulation of EC weights) within and between these maps. To examine this hypothesis, we implement a data-driven model of EC informed by the empirical temporal autocovariance of the BOLD data. This model reproduces the spatiotemporal covariance structure (the zero-lag covariance and time-lag covariance) of the empirical data and the local cortical excitability. Our model thus accounts for the spatiotemporal statistics of BOLD activity and thereby the propagation of BOLD activity across different brain locations [21, 20]. We apply this approach to resting state (RS) and visual field mapping (VFM) fMRI recordings of the early cortical visual field maps V1, V2 and V3 in healthy human participants.

In both RS and VFM data, we find a common structure underlying the EC of all participants, regardless of inter-participant variation in EC estimates. The common structure in EC links homotopic regions with similar visual field position selectivity both within and between the early cortical visual field maps, i.e. across both their topography and hierarchy. Furthermore, the resulting ECs capture different interaction regimes in RS and VFM. Within area interactions are increased in RS, whereas in VFM between area interactions are increased, particularly feedback interactions from V3 to V1. Moreover, local cortical excitability in V1 is greatly increased during RS but, during VFM, decreases to levels that are comparable to that of other visual areas. These differences point to a change of input to V1, and appear to reflect a re-configuration of feedforward, lateral and feedback interactions. Finally, we interpret our results under the framework of predictive coding, emphasizing the role of recurrent cortical feedback during visual processing. Taken together, our results demonstrate that the propagation of BOLD activity through early visual cortex, as assessed by the present analysis, have functional relevance.

## 2 Methods

### 2.1 Data

The empirical data used in this study stem from the dataset presented in Gravel et al. (2014) [23]. It comprise visual field mapping (VFM) and resting state (RS) 7T fMRI data from four healthy human participants (2 females, 2 males, age 26-40) with normal visual acuity. Experimental procedures were approved by the medical ethics committee of the University Medical Center Utrecht.

#### 2.1.1 *Visual field mapping*

Visual stimuli were presented by back-projection onto a 15.0 × 7.9 cm gamma-corrected screen inside the MRI bore. Participants viewed the display through prisms and mirrors, and the total distance from the participants eyes (in the scanner) to the display screen was 36 cm. Visible display resolution was 1024×538 pixels. The stimuli were generated in Matlab (Mathworks, Natick, MA, USA) using the PsychToolbox [8, 66]. The visual field mapping paradigm consisted of drifting bar apertures at various orientations, which exposed a 100% contrast checkerboard moving parallel to the bar orientation. After each horizontal or vertical bar orientation pass, 30 s of mean-luminance stimulus were displayed. Throughout the VFM, participants fixated a dot in the center of the visual stimulus. The dot changed color between red and green at random intervals. To ensure attention was maintained, participants pressed a button on a response box every time the color changed. Detailed procedures can be found in [14] and [30]. The radius of the stimulation area covered 6.25 deg (eccentricity) of visual angle from the fixation point.

#### 2.1.2 *Resting state*

During the resting state scans, the stimulus was replaced with a black screen and participants closed their eyes. The lights in the scanning room were off and blackout blinds removed light from outside the room. The room was in complete darkness. Thus, visual stimulation was minimized. The participants were instructed to think of nothing in particular without falling asleep.

#### 2.1.3 *fMRI acquisition*

Functional T2*-weighted 2D echo planar images were acquired on a 7 Tesla scanner (Philips, Best, Netherlands) using a 32 channel head coil at a voxel resolution of 1.98 × 1.98 × 2.00, with a field of view of 190 × 190 × 50 mm. TR was 1500 ms, TE was 25 ms, and flip angle was set to 80°. The volume orientation was approximately perpendicular to the calcarine sulcus. In total, eight 240 volumes functional scans were acquired, comprising 5 resting state scans (RS) interleaved with 3 VFM scans (first was an RS scan). Five dummy volumes were scanned before data acquisition began and a further eight volumes were discarded from the beginning of each scan to ensure the signal had reached a steady state. High resolution T1-weighted structural images were acquired at a resolution of 0.49 × 0.49 × 0.80 mm (1 mm isotropic resolution for the second dataset), with a field of view of 252 × 252 × 190 mm. TR was 7 ms, TE was 2.84 ms, and flip angle was 8°. We compensated for intensity gradients across the image using an MP2RAGE sequence, dividing the T1 by a co-acquired proton density scan of the same resolution, with a TR of 5.8 ms, TE was 2.84 ms, and flip angle was 1°. Physiological recordings were not collected.

#### 2.1.4 *Preprocessing*

First, the T1-weighted structural volumes were resampled to 1 mm isotropic voxel resolution. Gray and white matter were automatically labeled using Freesurfer and labels were manually edited in ITKGray to minimize segmentation errors [80]. The cortical surface was reconstructed at the white/gray matter boundary and rendered as a smoothed 3D mesh [86]. Motion correction within and between scans was applied for the VFM and the RS scans [61]. Subsequently, data were aligned to the anatomical scans and interpolated to the anatomical segmentation space [61]. Instrumental drift was removed by detrending with a discrete cosine transform (DCT) filter with cutoff frequency of 0.01 Hz. The detrended signals were used for the estimation of the empirical spatiotemporal covariances. In order to reduce the influence of high frequency variation during pRF modeling, the detrended signals were filtered with a low-pass 4th order Butterworth filter with cutoff frequency of 0.1 Hz.

### 2.2 Selection of regions of interest

Since the focus of our study was modeling the propagation of BOLD activity within and between early visual field maps, we did not consider all recorded locations in the scanning volume. Instead, we applied an ROI selection and a data-reduction step. First, we identified the visual field maps of the visual cortical areas V1, V2 and V3 (section 2.2.1). Second, we averaged the BOLD signals over the foveal and peripheral quarter-fields of these maps. This resulted in a network of 24 nodes/ROIs per participant (section 2.2.2).

#### 2.2.1 *Population receptive field modeling*

The visual field maps of V1, V2, and V3 were obtained using the population receptive field (pRF) method [14] applied to our VFM data. This method provides models that summarize the visual field position to which each recording site responds as a circular Gaussian in visual space. The Gaussian pRF model for each recording site was characterized by three parameters: x and y (position), and size (sigma). These parameters were determined by taking a large set of candidate pRF parameters, with each set defining a different Gaussian. By quantifying the overlap between each candidate pRF Gaussian and the stimulus aperture at each time point, we generate predictions of the neuronal response time course each candidate pRF would produce. This predicted neuronal response time course is convolved with the hemodynamic response function (HRF) to give a set of candidate predicted fMRI response time courses for each candidate set of pRF parameters. The best fitting predicted fMRI time course and its associated pRF parameters are then taken to summarize the visual field selectivity of each recording site [14]. Recording sites were excluded from subsequent analyses if their best fitting pRF models explained less than 30% of response variance, or had visual field eccentricities beyond 6 deg.

#### 2.2.2 *Grouping of data into foveal and peripheral quarter-fields*

We grouped the resting state (RS) and the visual field mapping (VFM) time series over the foveal and peripheral quarter-fields of V1, V2 and V3 using the eccentricity and polar angle pRF preferences of each recording site. The foveal ROIs grouped recording sites with pRF positions below 2.2 deg eccentricity, while peripheral ROIs grouped recording sites with pRF positions above 2.2 deg eccentricity. Quarter-fields were divided at the vertical and horizontal visual field meridians. The grouping process resulted in a matrix of 24 nodes/ROIs, 8 for each complete visual field map (V1, V2 and V3). Signals for each of the 24 ROIs were obtained by averaging the minimally preprocessed BOLD time series (after detrending) within the ROIs.

### 2.3 Effective connectivity model for BOLD propagation

In this section we examine the propagation of BOLD activity across the foveal and peripheral quarter-fields of V1, V2 and V3 (each of the 24 ROIs previously defined) using a recently proposed method [21]. This approach uses a noise-diffusion network model of effective connectivity (EC) and intrinsic variability to account for local BOLD variability and signal propagation lags between all possible pairs of ROIs. Importantly, the model captures the empirical data covariance and its spatio-*temporal* structure (the time-shifted covariances), effectively accounting for the propagation of BOLD activity. This has the advantage of relying on minimal assumptions: 1) the time constant of the generative model has to match the autocovariance time constant derived empirically from the data, 2) the regional variation in the hemodynamic response shape across early visual cortex should be minimal [26, 48] and 3) for each behavioral condition, a dominant pattern of neuronal interactions should influence and reflect in the average propagation structure of the BOLD signals. To examine our hypothesis (BOLD activity propagation reflects the consequences of structured neuronal activity), we model the EC under two conditions: 1) resting state and 2) the presentation of visual field mapping stimulus, and compare the two. We iteratively tune the model parameters (directed connectivity with notation *C* and intrinsic variability with notation *Σ* and) to reproduce the empirical spatiotemporal covariance and then use the *C* and *Σ* associated with the best fitting model as an estimate of the EC and the local cortical excitability of the actual data. We conclude comparing the resulting differences in EC and cortical excitability between RS and VFM.

#### 2.3.1 *Empirical spatiotemporal covariances*

To identify the spatiotemporal covariance structure of the data (the BOLD signals from each of the 24 ROIs. see Methods section 2.2) we estimated the covariance with and without time-shifts. For each participant and condition, the BOLD signals were first demeaned and then, following Gilson et al. (2016) [21], the empirical covariance was calculated for zero-lag:

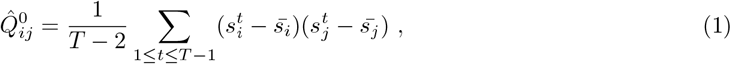

and a lag of 1 TR:

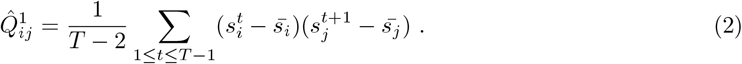

Here 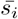 is the mean BOLD level over the session for ROI *i*. Afterwards, for each participant and session, we estimated the empirical time constant associated with the exponential decay of the autocovariance (averaged over all regions):

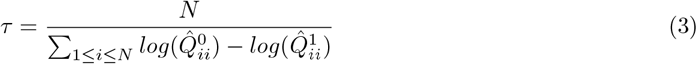

The time constant was used to calibrate the noise diffusion network model.

#### 2.3.2 *Noise-diffusion network model of EC and parameter estimation*

We choose a dynamic network model that captures the spatiotemporal dynamics of the data. Here we summarize the essential ingredients of the model and its optimization (for further details see Gilson et al. (2016) [21]). The models consists of 24 interconnected nodes (as defined in Methods section 2.2) that experience fluctuating activity and excite each other [21]. The local variability is described for each node by a variance corresponding to a diagonal term of the the matrix *Σ*. The implicated fluctuations are shaped by the network EC (denoted by the matrix *C* in the following equations) to generate the model FC, which is quantified the the zero-lag covariance matrix *Q*^0^ (FC0) and the time-lag covariance matrix *Q*^1^ (FC1) (the counterparts of the empirical 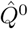 and 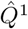). Subsequently, the model covariance matrices *Q*^0^ and *Q*^1^ that better reproduce the empirical spatiotemporal covariances 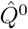 and 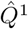 are approximated by iteratively adjusting the directional weights (*C*) and node excitabilities (*Σ*) of the model using Lyapunov optimization (LO) to reduce the model error *E*. The parameters *C* and *Σ* associated with the best fitting model correspond to maximum-likelihood estimates [21]. Importantly, because asymmetry in the *C*_*ij*_ generates asymmetry in the time-shifted covariances *Q*^1^, the model captures the average propagation structure between ROIs.

We choose LO because it has several advantages to other methods [21]: 1) pairwise unconditional Granger causality does not take the whole network into account; 2) multivariate autoregressive models (MVAR) models that take the whole network into account may suffer from the down-sampling due to the time resolution (TR = 1.5 s); 3) physical interpretability might be hindered by over-parameterized dynamic causal models (DCM) [22, 27, 78]. These advantages allowed us to estimate *C*_*ij*_ (and the corresponding asymmetry in *Q*^1^) and *Σ* as accurately as possible. Our approach was also justified because the decay time constant *τ* in Eq. (3)) was consistently measured across participants, suggesting a diffusion process in the empirical data; the goal of our model inversion was then to exmine whether propagation was present in the data.

Now we detail the equations relating these parameters, observables and measures. Formally, the network model is a multivariate Ornstein-Uhlenbeck process where the activity *x*_*i*_ of node *i* decays exponentially with the time constant *τ* estimated from the data in Eq. (3). The evolution of each *x*_*i*_ depends on the activity of other populations and the local variability:

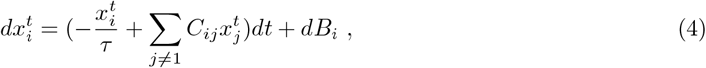
where *dB*_*i*_ is - both spatially and temporally - independent Gaussian noise with variance *Σ*_*ii*_ (the *Σ* matrix is diagonal); formally *B*_*i*_ a Wiener process. The model *Q*^0^ can be calculated for known *C* and *Σ* by solving the Lyapunov equation (using the Bartels-Stewart algorithm):

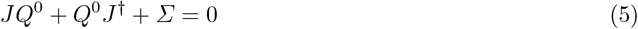
and *Q*^1^ is then given by

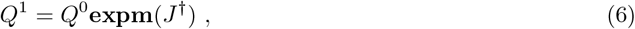

where **expm** denotes the matrix exponential, the superscript † indicates the matrix transpose, and *δ*_*ij*_ is the Kronecker delta. In those equations, the Jacobian *J* of the dynamic system is defined as

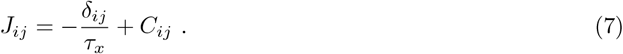

Eqs. (5) and (6) enable the quick calculation of *Q*^0^ and *Q*^1^, without simulating the network activity.

The Lyapunov optimization (LO) starts with zero connectivity (*C* = 0) and uniform local variances (*Σ*_*i*_*i* = 1). Each iteration of LO aims to reduce the model error defined as

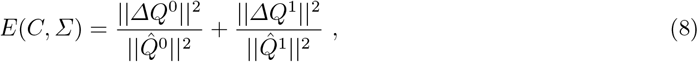
with the difference matrices 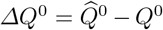 and 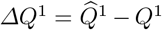; ‖ · ‖ indicate the usual matrix norm. To do so, we calculate the model *Q*^0^ and *Q*^1^ for the current values of the parameters *C* and *Σ* by solving Eqs. (5) and (6). Similar to a gradient descent, the Jacobian update is given by:

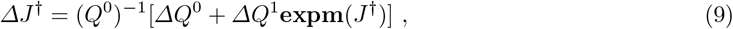
which gives the connectivity update:

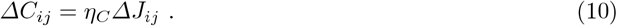
Where *η*_*C*_ is the optimization rate of *C* (here we *η*_*C*_ = 0.0001). To take properly the network effects in the *Σ* update, we adjust the *Σ* update from the heuristic update in [21] as was done in [20]:

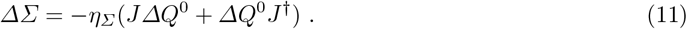
Where *η*_*Σ*_ is the optimization rate of *Σ* (here *η*_*Σ*_ = 1). We impose non-negativity both for *C* and *Σ*. In addition, off-diagonal elements of *Σ* are kept equal to 0 at all times. The optimization steps are repeated until reaching a minimum for the model error *E*, giving the best fit and the model estimates. The *C* and *Σ* associated with best predicting model are then taken as a proxy for the EC between the 24 ROIs and their local cortical excitability (*Σ*). To facilitate comparison between RS and VFM, the resulting *Σ* values were further grouped across foveal and peripheral ROIs in each hemisphere (giving a total of 6 new ROIs) and differences between RS- and VFM-derived values evaluated for significance using single sided permutations. A similar approach was applied to the resulting *C*_*ij*_ values (see next section). Further details about the mathematical formalism of the model and the optimization procedures can be found in Gilson et al. (2016 & 2017) [21, 20].

#### 2.3.3 *Determination of common underlying structure in EC and its relation to topographic and anatomical connectivity*

Due to the nature of the model parameter estimation, the estimated EC weights (gain) may incidentally vary across participants (i.e. due to inter-participant variation, data quality, etc). However, relative differences in EC gain may still be similar across participants, suggesting a common underlying structure. To identify if there was a common structure underlying the EC of all participants, raw EC estimates were first normalized by correcting for relative changes in EC gain across participants. Importantly, this process renders individual EC estimates comparable (since weight magnitudes are normalized) and enables the computation of an unbiased grand average. Gain correction was possible because, regardless of individual variation in EC gain across participants, the comparison of raw EC gain values between each of the 6 possible participant pairs revealed correlated slopes (The Pearson correlation coefficient between individuals EC over the 6 possible combinations was, in RS: 0.28, 0.48, 0.43, 0.23, 0.23, 0.37 with *p* < 10^−7^ in all cases, and, for VFM: 0.67, 0.73, 0.71, 0.58, 0.61, 0.68 with *p* < 10^−54^ in all cases), which we interpret as similar relative pairwise differences between EC links (and therefore similar structure).

To normalizes EC weights across participants, we re-scaled the individual EC values by realigning the slopes (estimated as the Pearson correlation coefficient) of each pairwise EC difference to 1(based on an arbitrary reference participant). Subsequently, the normalized EC matrices were classified as intra- or inter-hemispheric and normalized EC links that appeared consistently across participants were detected using permutations corrected for multiple comparisons (with a threshold of *p* < 0.05). This allowed us to detect the common structure underlying the EC of all participants representing it as a binary matrix of significant connections. We then went on to examine differences in EC between RS and VFM. To this end, the normalized EC values that matched the matrix of significant connections were averaged across participants resulting in a grand average EC matrix. The averaged EC values were grouped across all quarter fields to give smaller EC matrices consisting of 6 ROIs (foveal and peripheral parts of V1, V2 and V3). Changes in EC between conditions were then quantified by computing the difference between the resulting VFM- and RS-derived grand average EC matrices. Finally, differences between RS and VFM were evaluated for significance across participants using permutations corrected for multiple comparisons.

## 3 Results

### 3.1 Propagation of BOLD activity across early visual cortex measured with a noise-difussion network model of EC

Figure 1 illustrates the propagation of an apparent wave of BOLD activity from the anterior calcarine sulcus (periphery of V1) to the occipital pole (foveal confluence of V1, V2 and V3) during rest (RS) (see Video 1S in supplementary materials). To estimate the propagation of BOLD activity across early visual cortex we first obtained visual field maps of V1, V2 and V3 using the population receptive field (pRF) method (Fig 2A). We then further subdivided these maps into foveal (below 2.2 deg of eccentricity) and peripheral (above 2.2 deg of eccentricity) quarter-fields (Methods section 2.2). This provided us with a functional map of cortex based on similarities in both retinotopy and hierarchy. Second, we characterized BOLD activity propagation patterns during RS and VFM through V1, V2 and V3 using a data-driven modeling approach (Methods section 2.3). Here we used the temporal autocovariance constant derived empirically from the data to calibrate a topologically agnostic (unconstrained by anatomical connections) noise diffusion network model of EC and cortical excitability. Figure 2B presents the results from the analysis of the temporal autocovariance. Different time constants (*τ*, in seconds) for RS and VFM (mean (std) = 3.64 (1.46)) for RS and 6.66 (0.98) for VFM, paired t-test: *p* < 10^−5^) demonstrate different propagation regimes. We then modeled the spatiotemporal covariance structure of the data by optimizing the noise-diffusion network parameters, namely the effective connectivity (EC) and the nodes excitabilities (*Σ*), to reproduce the empirical spatiotemporal covariances FC0 and FC1. Figure 2C illustrates one iteration step in the Lyapunov optimization (LO) procedure used to solve the model. The goodness o fit between modeled and empirical spatiotemporal covariances was computed using the linear regression coefficient *R*^2^ between the modeled and the empirical FC0 and FC1 (see Table 1).

**Fig. 1:**
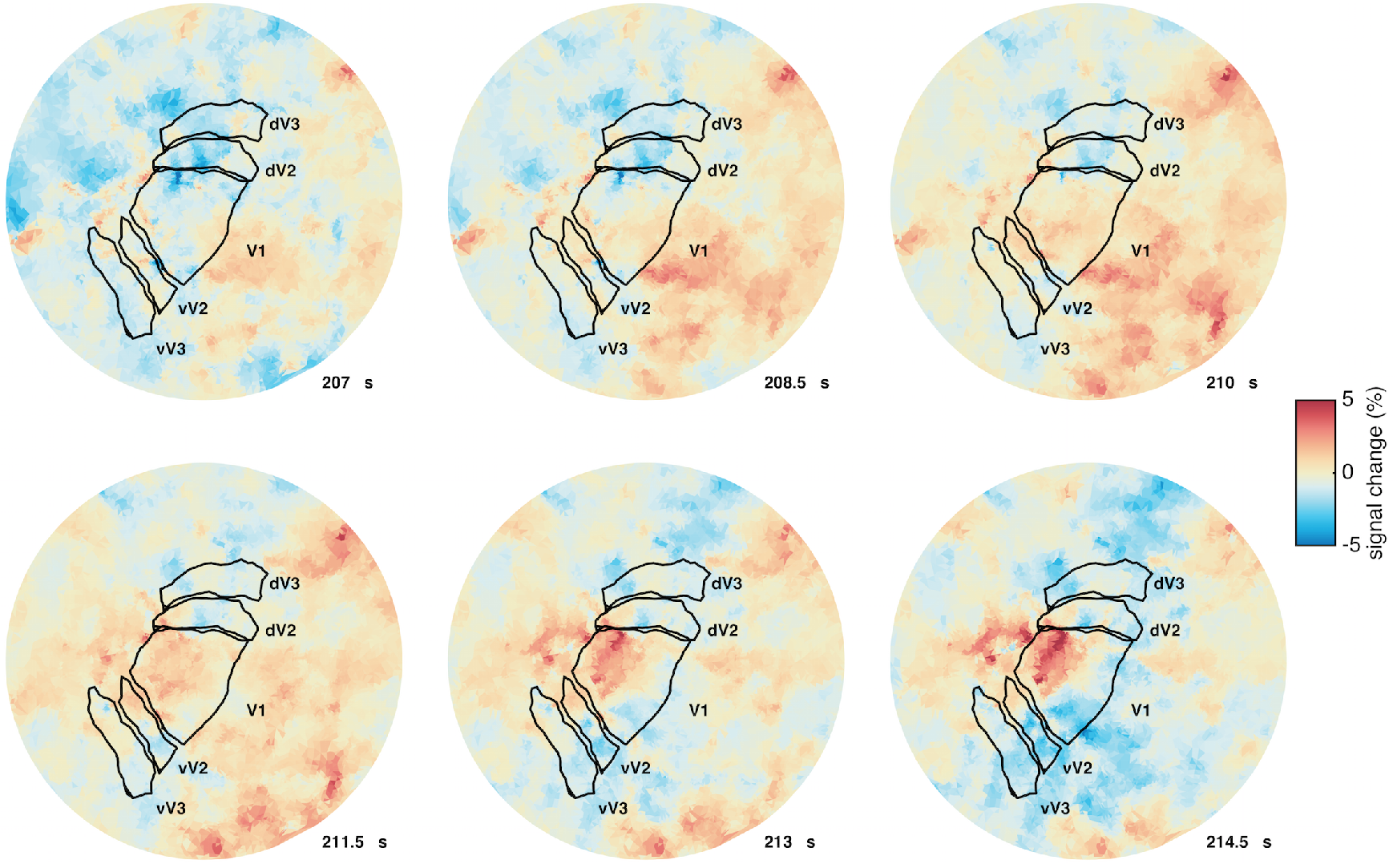
Apparent propagation of BOLD activity during RS depicted in the flattened cortical surface reconstruction of the occipital pole of one participant's cerebral hemisphere. Early visual field maps V1, V2 and V3 in one hemisphere are outlined in black (d- and v- denote dorsal and ventral). In the absence of visual stimulation (i.e. eyes closed, total darkness), spontaneous fluctuations in BOLD activity (indicated by the hot and cold colors) during RS can exhibit extensive spatiotemporal structure. This structure includes transient spatiotemporal fluctuation patterns that resemble stimulus-evoked BOLD waves as well as congruent and transient co-activations that occur across the visual field maps. Video 1S in supplementary materials shows a movie illustrating this propagation of BOLD activity.

**Fig. 2:**
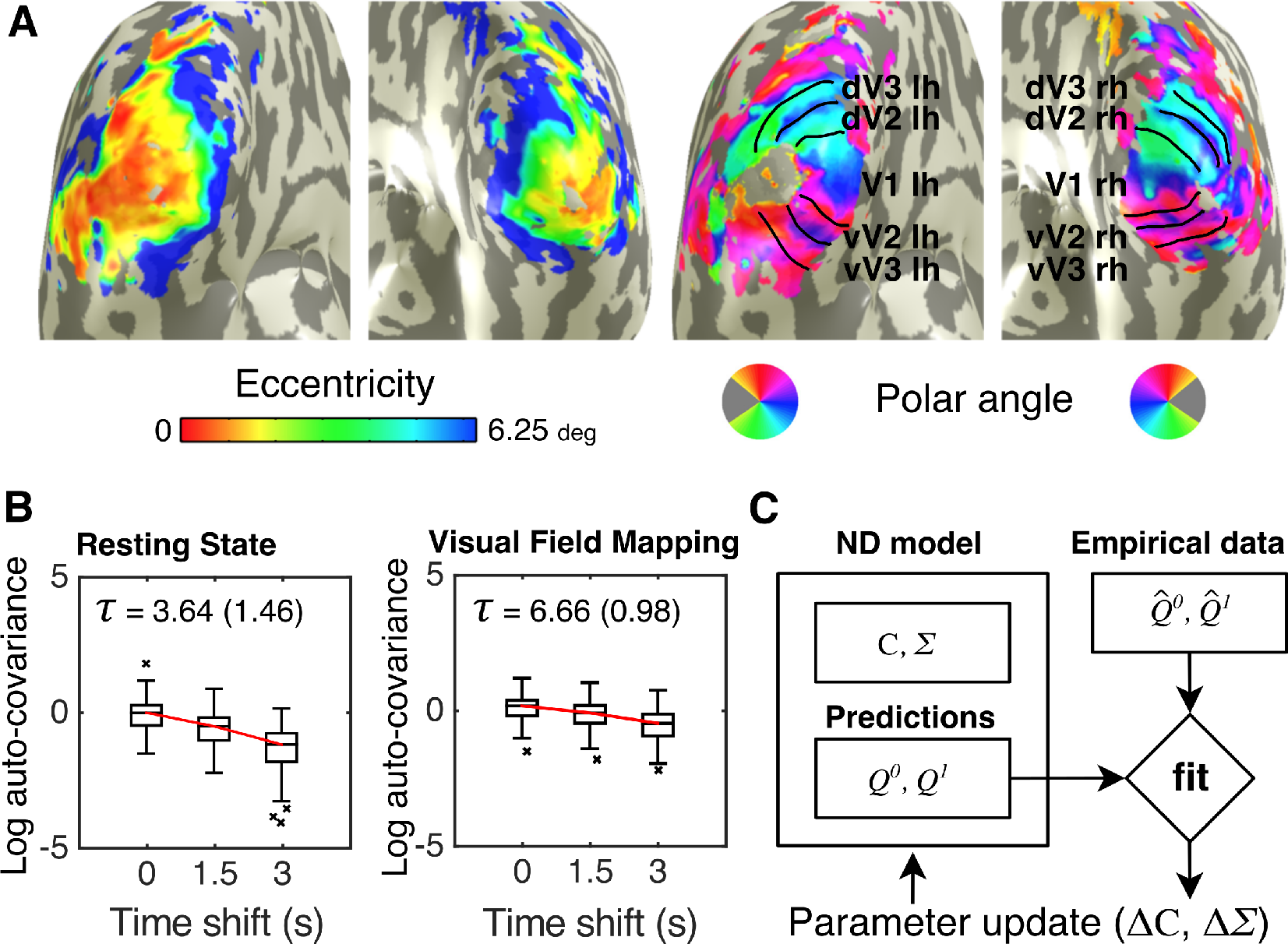
Modeling the propagation of BOLD activity across visual field maps V1, V2 and V3. **(A)** Visual field maps in striate (V1) and extrastriate cortex (V2 and V3) were mapped using the population receptive field (pRF) modeling method [14]. Based on the visual field position selectivity estimates that the pRF method provides, these maps were further subdivided into foveal (below 2.2 deg of eccentricity) and peripheral (above 2.2 deg of eccentricity) quarter-fields in both hemispheres, giving a total of 24 ROIs (Methods section 2.2). We used a noise diffusion network model of effective connectivity (EC) to estimate the propagation of BOLD activity and the cortical excitability across these foveal and peripheral quarter-fields of V1, V2 and V3. **(B)** Logarithm of the autocovariance of the BOLD activity, averaged across participants and ROIs, as a function of the time shift (x-axis). The red lines link the mean of each box plot. The time constant *τ* was calculated for each participant and condition (Methods section 2.3.1). The mean (standard deviation) of *τ* over all participants is indicated above the box plots. The empirical time constants were used to calibrate the noise-diffusion network model. **(C)** Schematic diagram illustrating one step in the Lyapunov optimization (LO) procedure. By iteratively adjusting the connectivity C and node excitability *Σ* of the noise-diffusion network, the model spatiotemporal covariance (Q^0^, Q^1^) approximates the empirical spatiotemporal covariance (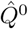,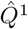). For each participant and condition, the *C* and *Σ* corresponding to the best predicting model were taken as estimates of the underlying effective connectivity (EC) and the local cortical excitability (model optimization results for all participants are given in supplementary Figure 1S).

**Table 1:** Goodness of fit between modeled and empirical spatiotemporal covariances. For each participant and condition, we evaluated the goodness of fit by computing the linear regression (*R*^2^, *p* < 10^−50^ for all cases) between the modeled and the empirical spatiotemporal covariances (FC0 and FC1).

| participant | RS |  | VFM |  |
| --- | --- | --- | --- | --- |
|  | FC0 | FC1 | FC0 | FC1 |
| 1 | 0.652 | 0.629 | 0.694 | 0.655 |
| 2 | 0.473 | 0.329 | 0.702 | 0.689 |
| 3 | 0.884 | 0.829 | 0.705 | 0.665 |
| 4 | 0.847 | 0.819 | 0.754 | 0.731 |
| **Mean (std):** | 0.71 (0.19) | 0.65 (0.23) | 0.71 (0.027) | 0.68 (0.033) |

### 3.2 Common underlying structures in EC

We then asked if there was a common structure underlying the resulting ECs. Regardless of individual variations in EC values obtained from RS and VFM data (Fig 3A,E), a common underlying structure was revealed both for RS and VFM. Figure 3B and 3F compare relative pairwise differences between EC links in different participants pairs. The Pearson correlation coefficient between individuals EC over the 6 possible combinations was, in RS: 0.28, 0.48, 0.43, 0.23, 0.23, 0.37 with *p* < 10^−7^ in all cases, and, for VFM: 0.67, 0.73, 0.71, 0.58, 0.61, 0.68 with *p* < 10^−54^ in all cases. These results indicate that similar relative pairwise differences between EC links (and therefore similar structure). Figure 3C and 3G compare the normalized ECs to the grand average EC. Despite individual variation in overall EC gain, the normalized ECs for all participants were highly correlated to the grand average EC. The linear correlation coefficient *R*^2^ between the normalized individual ECs and the grand average EC was: 0.60, 0.45, 0.46 and 0.47 for RS and 0.81, 0.67, 0.76 and 0.74 for VFM (*p* < 10^−50^ in all cases). These results indicate that differences between individual ECs are less pronounced in VFM. Figure 3D and 3H depict the EC links that appeared consistently across participants as directed binary matrices. We used these matrices as estimates of the common structure underlying the EC in RS and VFM. The common structures in EC closely matched the homotopy and hierarchy of the underlying anatomical connections (The Pearson correlation coefficient between the EC-structures and binary matrices for within-hemisphere and between-hemisphere connections with ones indicating homotopy was: 0.52 for VFM with *p* < 10^−40^ and 0.24 for RS with *p* < 10^−9^). However, the homotopic organization of these was better captured in VFM (Fig 3H).

**Fig. 3:**
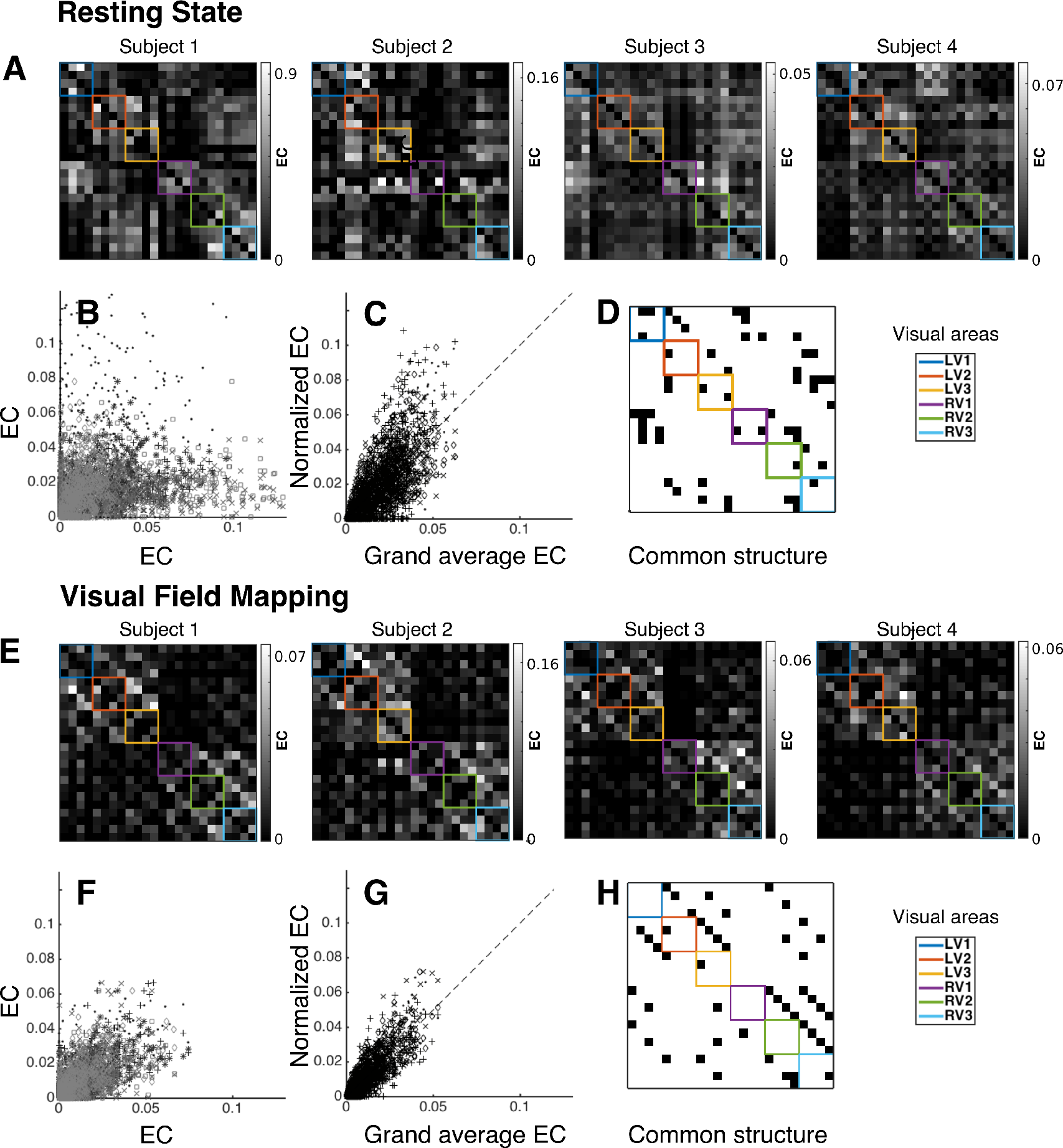
Common underlying structure of EC across visual cortical areas V1, V2 and V3. **(A)** Diagonal and off-diagonal quadrants in each matrix represent within- and between-hemisphere EC across visual cortical areas (grouped by the colors), respectively. Inside each colored box, quarter-fields are grouped in the following order (from left to right): upper fovea, upper periphery, lower fovea and lower periphery. For each diagonal and off-diagonal quadrant, the upper triangle represents feedback connections and the lower triangle feedforward connections (rows correspond to inputs and columns correspond to outputs). White pixels represents stronger EC weights and darker pixels weaker EC weights, as indicated by the colorbars. **(B)** Scatter-plot comparing raw EC values between different participants. Different markers depict each of the 6 possible pairs. **(C)** Scatter-plot comparing the normalized EC values of each participant to the grand average EC. Different markers depict each participant. Despite individual variation in overall gain, the normalized EC for all participants was highly correlated to the grand average. **(D)** Common structure in EC. Black dots indicates connections that appeared consistently across participants. Significance was evaluated using permutations corrected for multiple comparisons (*p* < 0.05). **(E-H)** Same as **A-D** but for VFM. Differences between individual ECs were less pronounced for VFM data.

### 3.3 Differences in EC and *Σ* between RS and VFM

We then went on to examine the topology of the resulting common structures in EC and their differences between conditions. Figure 4 illustrates the common structures in EC (significant intra- and inter-hemispheric connections) across the foveal and peripheral quarter-fields of the visual field maps for RS and VFM. Both in RS and VFM, intra- and inter hemispheric connections linked regions with similar visual field selectivity, however, less inter-hemispheric connections were detected. Interestingly, in VFM, inter-hemispheric connections linked foveal regions only, whereas, in RS inter-hemispheric connections linked peripheral regions only. Both in RS and VFM, feedback connections outnumbered feedforward connections.

**Fig. 4:**
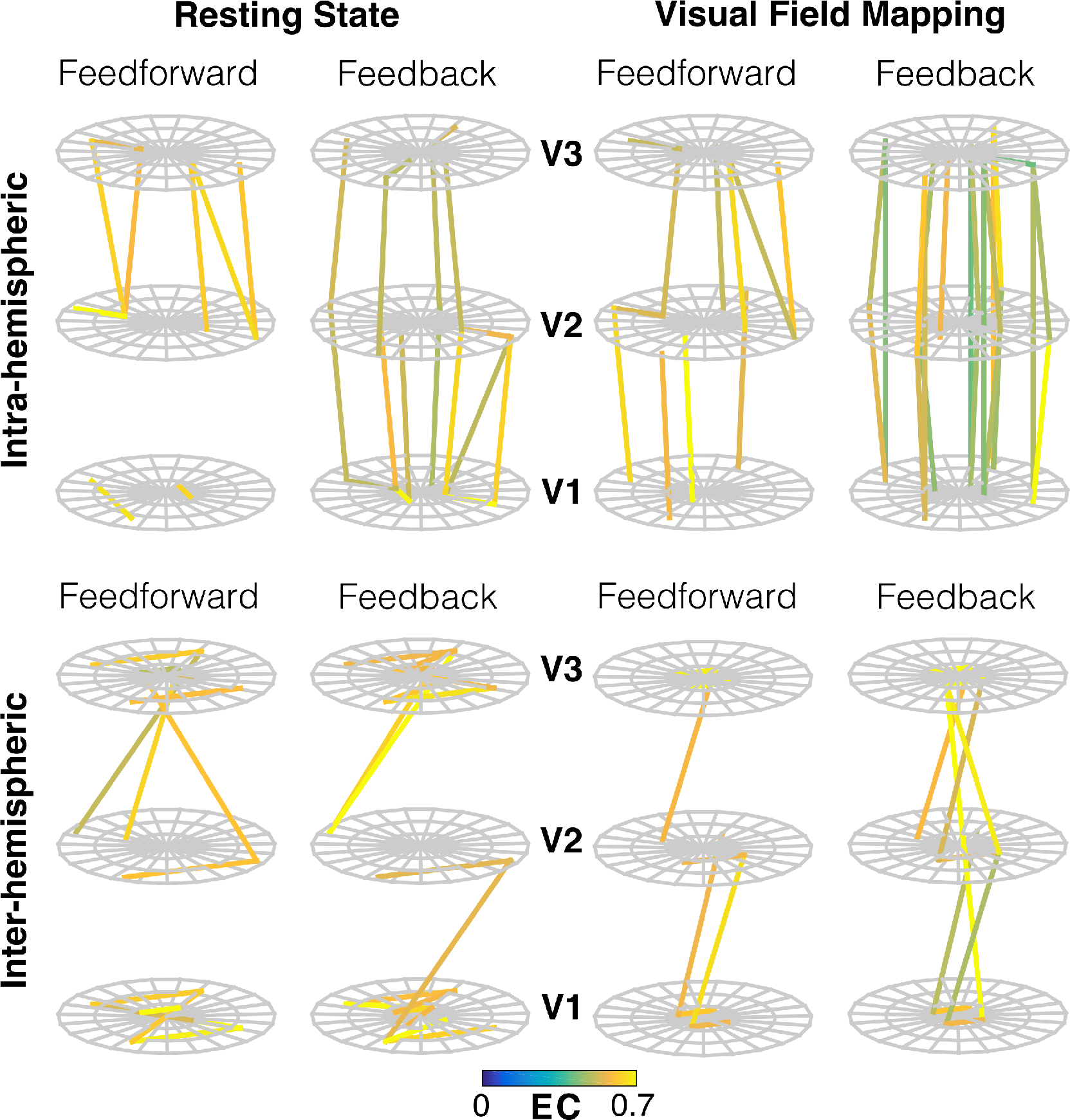
EC structure for RS and VFM illustrated in visual field space. The common structures in EC from Figure 3 (D,H) are displayed here as networks overlaying 3D representations of the visual field position selectivity of the 24 ROIs defined in section 2.2 (vertically stacked grids) by separating: 1) intra- and inter-hemispheric links, and 2) feedforward and feedback links. Colors depict the strengths of the grand average ECs (see Methods section 2.3.3 for details). In order to avoid overlaps between V1-V3 and V1-V2/V2-V3 links, we shifted the eccentricity associated to the visual position selectivity grid of V2 slightly to the periphery. The EC networks thus depicted suggest different interaction regimes in RS and VFM (see Figure 5 for a detailed assessment of these differences). The vertically stacked grids are oriented to match the left/right orientation of the visual field.

Subsequently, to interpret changes in the EC and cortical excitability between RS and VFM, we grouped the corresponding EC and *Σ* values across the four quadrants in each visual field map to give foveal and peripheral regions of each visual field map (the 6 ROIs defined in section 2.3.3). We then evaluated differences in EC between the two conditions across participants using permutations corrected for multiple comparisons (with a significance threshold of *p* < 0.05). Figure 5A illustrates the resulting ECs and the differences between conditions (VFM-derived EC minus RS-derived EC). These results show that strong interactions within V1 and V3 in RS are absent in VFM. Conversely, feedforward interactions between V1 and V2 were present only in VFM, whereas feedforward interactions between V2 and V3 were present both in RS and VFM, although foveal interactions were increased for VFM. Further, feedback interactions between V2 and V1 and V3 and V2 were present both in RS and VFM, although feedback interactions between V3 fovea and V2 fovea were greatly increased in VFM. Notably, homotopic feedback interactions between V3 and V1 were only detected in VFM. Figure 5B illustrates the differences in cortical excitability between RS and VFM, as estimated by *Σ*. Changes were most pronounced in V1, with higher values of *Σ* for RS. Cortical excitability was significantly higher for foveal regions of V1 only (*p* = 0.033. Single sided permutation test).

**Fig. 5:**
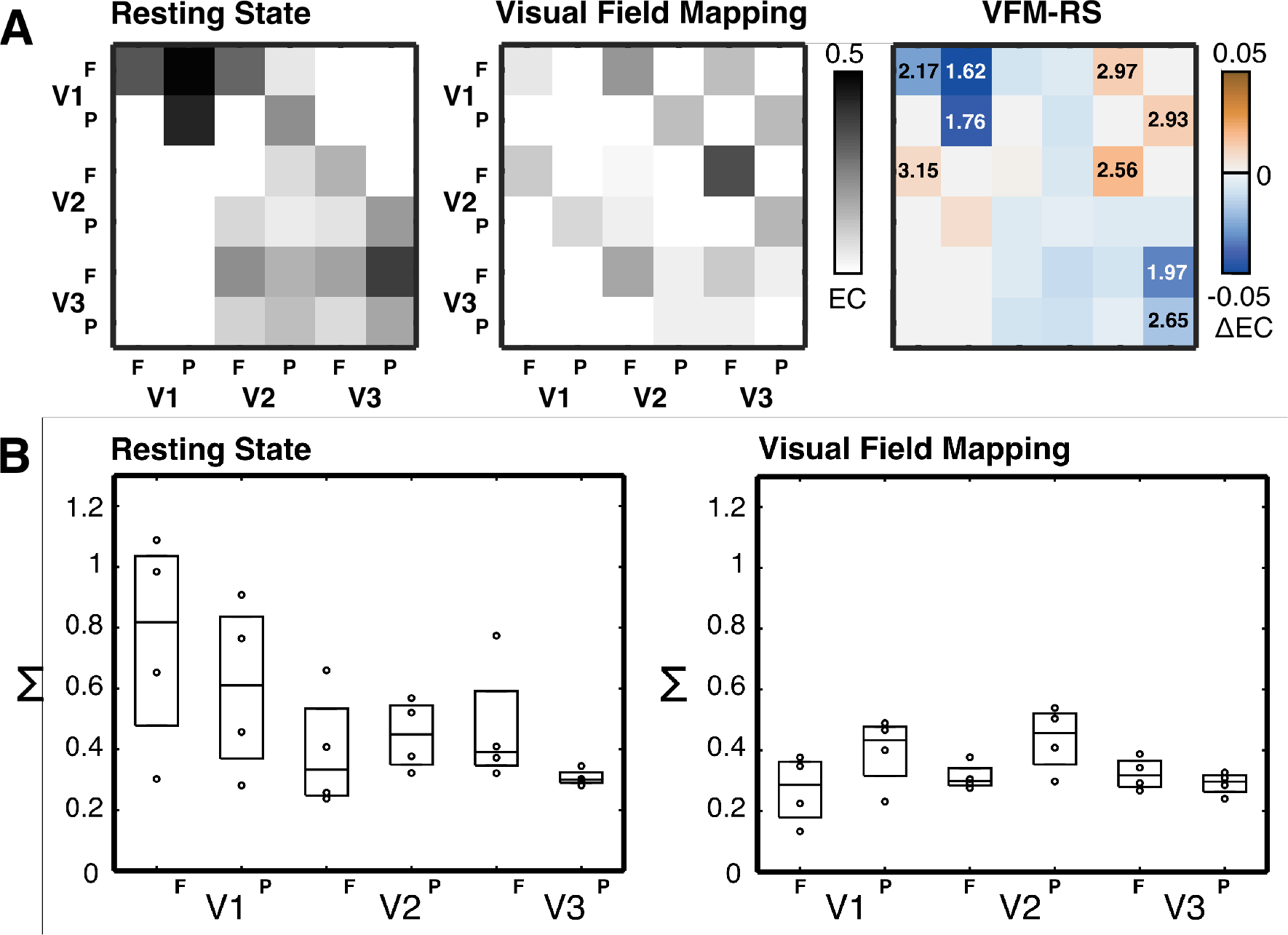
Differences in EC and *Σ* between RS and VFM suggest a re-configuration of feedforward and feedback interactions. **(A)** Average EC matrices for RS (left panel) and VFM (middle panel) obtained by grouping significant EC links into foveal and peripheral regions V1, V2 and V3 (F and P stand for fovea and periphery. Columns represents output and rows inputs. See Methods section 2.3.3 for details). The right panel illustrates the differences in EC between conditions (VFM-derived EC minus RS-derived EC). Cells corresponding to regions that showed significant changes in EC have annotated the negative logarithm of their p-value (−log_10_(*p*), note that − 1og_10_(0.05) = 1.3). **(B)** Cortical excitability as estimated by *Σ*. For each condition, individual *Σ* estimates were grouped into foveal (F) and peripheral (P) regions (as in A) and represented as black dots overlaid on box-plots (the central mark is the median and the edges the 25th and 75th percentiles). A comparison between RS and VFM reveal a slight decrease in this value, particularly for V1 (*p* = 0.033 for V1 fovea, *p* = 0.118 for V1 periphery, *p* = 0.33 for V2 fovea, *p* = 0.46 for V2 periphery, *p* = 0.08 for V3 fovea, and *p* = 0.271 for V3 periphery. Single sided permutation test). We believe that the increased values of *Σ* for V1 found in RS may relate to alpha activity (see Discussion section 4.2). In VFM, *Σ* is slightly increased in the periphery of V1 and V2 (although not significantly). Feedback connections outweighed feedforward connections both in resting state (RS) and visual field mapping (VFM).

## 4 Discussion

We assessed the propagation of BOLD activity through early visual cortical areas V1, V2 and V3 during RS and VFM using a data-driven modeling approach based on a noise-diffusion network model. Informed by the empirical spatiotemporal covariance structure of BOLD co-fluctuations within and between visual cortical areas, this model estimates a topologically agnostic (unconstrained by anatomical connections) effective connectivity (EC). Our model decomposes the spatiotemporal structure of BOLD fluctuations into an EC parameter and a local cortical excitability parameter *Σ*. Importantly, the combination of the estimated parameters explain the temporal lags between BOLD signals from all pairs of ROIs, effectively accounting for observed propagation in the data.

This discussion comprises four sections. In the first section we examine the neuroanatomical substrate and the possible mechanisms implicated by the different EC interactions estimated for RS and VFM. Our focus here is to emphasize the role of recurrent feedback connectivity and non-stimulus driven inputs, as well as examine the functional implications of our findings from a theoretical perspective—touching upon the notion of predictive coding [45, 69]. In the second section, we discuss the possible mechanisms that underlie the changes in cortical excitability (*Σ*) observed between RS and VFM and relate those to changes in EC. In the third section, we relate the BOLD autocovariance decay constant to different behavioral states. In the last section, we discuss the methodological and theoretical limitations of our study and raise questions for future research.

### 4.1 Recurrent connectivity and its role in visual processing

We demonstrate that the propagation of BOLD activity across the topography and hierarchy of (i.e. within and between) visual field maps V1, V2 and V3 reveals different directed interaction regimes for RS and VFM (Fig 3). We relate these differences in EC to a task-dependent reconfiguration of feedfoward and feedback interactions (Fig 4). Across visual field maps, feedforward EC interactions between the foveal representations of V1 and V2 were found in VFM but not in RS. However, later in the hierarchy, feedforward interactions between V2 and V3 were observed both in RS and VFM, though foveal interactions were increased in VFM (Fig 5A). We attribute these increased feedforward interactions during VFM, both between V1 and V2, and between V2 and V3, to stimulus-induced changes in neuronal interactions while participants are fixating on the screen, likely reflecting increased bottom-up processing of the stimulus across the visual hierarchy. Furthermore, homotopic feedback interactions between V2 and V1, and V3 and V2, were observed both in RS and VFM, although foveal interactions between V3 and V2 were greatly increased in VFM, again pointing to stimulus-induced changes. Remarkably, homotopic feedback interactions between V3 and V1 were only observed for VFM (Fig 5A), evidencing the role of extra-striate feedback in visual cortical processing.

At the level of individual visual field maps, we found directed EC interactions from the peripheral to the foveal representations of V1 and V3 in RS but not in VFM. In principle, these periphery-to-fovea interactions may reflect a bias in the spontaneous propagation of correlated neuronal activity along gradients of anatomically connected pathways or ‘connectopies’, such as the eccentricity map in the calcarine sulcus [24]. Also, other mechanisms such as intrinsic fluctuations in the ratio between excitation and inhibition and the modulation of lateral inhibitory coupling are known to give rise to spontaneous wave propagation patterns [15, 35]. Input changes, such as the restructuring of corticothalamic network activity, might adjust the balance between excitation and inhibition in cortical neuronal populations in a state-dependent way [79]. One interesting possibility is that these periphery-to-fovea interactions reflect large-scale cortical waves traveling in the frontal-to-occipital direction. Pre-stimulus waves in the alpha range [65] as well as slow <1 Hz waves propagating in an antero-posterior direction during sleep and calmness [52, 53] have been reported and may have a functional relevance [49, 37, 39]. In the absence of stimulation, the increased periphery-to-fovea EC interactions may well reflect visual cortical operations related to memory consolidation and learning [57]. On the other hand, the absence of periphery-to-fovea interactions during VFM can be explained by the stimulus set. Since the VFM stimuli consisted on a bar drifting at various orientations, the direction of neuronal response propagation may be balanced out and not reflected in EC estimates, which aim to assess stationary propagation patterns. Another line of evidence points to feedback modulation by top-down processes associated with the predictability of a given stimulus. Previous studies have suggested that predictable stimuli (e.g. drifting bars) induce less neuronal activity in early visual cortical areas than unpredictable stimuli [7, 28]. Concomitantly, this reduced neuronal activity induced by predictable stimuli has been shown to result in suppressed BOLD responses in early visual cortex [72, 73]. These results can also be interpreted within a predictive inference framework [85, 60, 45]. In this framework, recurrent feedforward/feedback loops serve to integrate top-down contextual priors (predictions) and bottom-up visual input by implementing a convergent probabilistic inference along the visual hierarchy. From this perspective, the increased feedback to V1 and V2 observed for VFM might be interpreted as the dampening of feedforward visual responses according to prediction errors originated by the mismatch between incoming signals and feedback priors [45, 69, 16, 18, 71].

### 4.2 Cortical excitability: possible mechanisms

Cortical excitability in RS, as quantified by *Σ*, was more variable than during VFM and was increased in V1, particularly in foveal representations (Fig 5B). These differences in cortical excitability between foveal and peripheral regions, although not consistently reaching statistical significance (*p* = 0.033 for V1 fovea), may still have functional implications. Interestingly, this increase in cortical excitability was accompanied by strong directed interactions in EC from the periphery to the fovea of V1 that were absent in VFM (Fig 5A). One possibility is that these differences are related to changes in the power of occipital alpha oscillations, known to increase during wakeful detachment from the environment (i.e RS) [87, 31]. If changes in the power of occipital alpha oscillations are partly captured by *Σ*, reduced cortical excitability in foveal regions of V1 could be interpreted as reflecting surround suppression and facilitation in the fovea while participants are fixating on the screen [31, 25]. Indeed, during a visual task, alpha oscillations in V1 have been associated with increased negative BOLD responses and shown to vary as a function of stimulus position and local receptive field surround [31]. The highly localized nature of these oscillations points to a role of intra-cortical axons in surround suppression [75, 74, 31, 36]. In line with these studies, changes in *Σ* were identified in the foveal representation of V1 but not in extrastriate areas V2 and V3, likely reflecting the localized nature of occipital alpha oscillations.

Extrastriate feedback to V1 may play a role modulating the balance between inhibition and excitation, known to change between task and rest, thereby reflecting differently in the BOLD signal during RS and VFM [44, 67]. In the absence of visual input (RS), changes in cortico-cortical connectivity may leave V1 in a ‘baseline’ state of potential excitation, whereas during VFM, both external visual input and top-down feedback influences may modulate the balance between inhibition and excitation resulting in suppressed BOLD responses in V1—thus attenuating cortical excitability [3, 72, 73]. The fact that we found lower values of *Σ* in the fovea of V1 for VFM corroborates this view.

### 4.3 Relation of the BOLD autocovariance decay constant to behavioral condition

The temporal decay constants of the autocovariance (*τ*), derived empirically from the RS and VFM data, determined the rate at which the fluctuations diffused through the noise-diffusion network model. Higher values of *τ* imply longer temporal memory (i.e. the systems past dynamics have a stronger influence on its future dynamics). Importantly, the determination of consistent different decay constants for RS and VFM demonstrated that propagation was present in both cases, albeit at different temporal scales. Estimates of *τ* obtained from VFM data were greater than those derived from RS data (Fig 2B), indicating longer temporal memory during VFM. This is in contrast to previous studies showing that temporal memory decreases during task compared to RS [32, 33]. By estimating task-induced decreases in the power-law exponent of BOLD fluctuations across widespread brain regions, these studies suggest that the temporal memory is longest during RS. They relate the larger power-law exponent found in RS to higher time-lagged autocovariances and interpret this as longer temporal memory. We find that the temporal memory (estimated by *τ* in our study) of visual cortical BOLD fluctuations is greater during task (here VFM) than RS. One possible reason for this differing results is that He and colleagues examined widespread whole-brain interactions, whereas here we examined BOLD signal dynamics at a more local scale (the cortical surface of individual visual field maps) and with higher resolution (7T). Furthermore, our results are specific to early visual cortex and therefore may not generalize to the whole brain. Another possible explanation is that these studies were based on eyes-open RS (fixation on a white cross-hair in the center of black screen) whereas we used eyes-closed RS. These different measurement scales and task protocols may well explain the observed differences.

In our study, the longer temporal memory found in VFM likely reflects stimulus induced interactions. By giving rise to slow frequency fluctuations in the BOLD signal that are spatially correlated with the stimulus position, these interactions may lead to higher temporal redundancy and therefore longer memory depth (higher *τ*). On the other hand, the absence of stimulus induced interactions during RS may lead to a decrease in spatiotemporally correlated slow fluctuations, leaving intrinsic fluctuations and fast transitions to dominate the temporal autocovariance structure (thus reducing *τ* during RS). Finally, the aforementioned studies [32, 33], computed the power-law exponent of the fMRI time series by using the low frequency range (< 0.1 Hz) of the power spectrum, whereas we computed *τ* from minimally preprocessed BOLD time series to which only detrending and demeaning was applied. By avoiding such low-pass filtering, we allowed faster fluctuations to influence our estimates of *τ*. All these lines of evidence suggest that, in early visual cortex, the temporal scale of BOLD activity propagation differs between RS and VFM. Compared to the slow and spatially widespread (long) propagation patterns evoked by the VFM stimulus, intrinsic fluctuations during RS tend to unfold locally in space and time. These shorter and more localized propagation events may dominate the spatiotemporal covariance structure and explain the increased EC within-area interactions observed during RS.

### 4.4 Limitations and interpretability of the model

An important limitation in the present study was the fact that we only considered a subset of all existing connections. This might have neglected the contribution of indirect connections to estimated changes in the EC, as dependencies may arise from indirect interactions in the underlying anatomy. Similarly, the increased *Σ* values identified during RS in the foveal representation of V1 may reflect input from other brain areas as well.

Other possible limitations derived from the fact that our approach was different from the original implementation of the noise-diffusion network model [21, 20] in two aspects: 1) Diffusion tensor imaging (DTI) can’t estimate structural connectivity at the spatial scales involved here. Therefore, our implementation was topologically agnostic compared to these previous studies: no structural connectivity matrix (i.e. DTI-derived) was used to constrain the EC; 2) we apply the approach to the scale of individual visual field maps whereas it was originally devised for whole-brain analyses (ROI ~ 500-1000 voxels instead of ~ 50 here) [21, 20]. However, we think the approach is still valid since there are known strong anatomical connections between V1, V2 and V3. Also, regional variation in the hemodynamic response is less pronounced at this scale [48], which further justifies the implementation of the model.

Another limitation is that we only allowed positive weights to be adjusted in the EC, which lead us to interpret the intrinsic variability of the model (*Σ*) to the aggregate changes in cortical excitability across both inhibitory and excitatory neuronal populations. We justify this decision based on the metabolic underpinnings of the BOLD signal: inhibitory functions, which are supported more by oxidative mechanisms than by excitatory signaling, may contribute less than excitatory functions to the measured BOLD activity [11]. Therefore, excitatory glutamatergic input to principal neurons might influence EC more than modulatory functions, which are exerted by a mostly inhibitory interneuronal network [11]. Furthermore, Allowing negative correlations between one voxel and another allows connections between a voxel where the stimulus is in the centre of the pRF and a voxel where the stimulus is in the suppressive surround. Supressive surrounds are large, so this is likely to lead to widespread spurious EC. Based on these reasons, we believe that positive weights in the EC are enough to capture the underlying neuronal interactions that shape BOLD responses.

We note that our aim was not to infer the detailed causal mechanism that give rise to the propagation of BOLD activity. Rather, our aim has been to assess the utility of a specific effective connectivity framework (a topologically agnostic noise-diffusion network) to quantify BOLD activity propagation at the level of the individual cortex. However, we note that local variation in neurovascular coupling profiles may hinder our analysis [4, 64], as they would affect the EC values (but less likely their modulations across conditions). Nevertheless, we do not model the hemodynamic response function (HRF) for a number of reasons. First, we assume the HRF to be relatively constant across V1, V2 and V3, even though the underlying vascular network may introduce certain non-uniformity [29, 26, 81, 48]. Second, our model reproduces the empirical spatiotemporal covariance analytically, and therefore does not rely on generative models of neuronal activity and neurovascular coupling to simulate the BOLD time series. While such an approach would be valuable to address questions of mechanistic causality, we think that the current temporal and spatial resolution of fMRI leaves such questions out of reach.

The time- and task-dependent nature of BOLD activity propagation patterns poses the question of how closely directed interactions map onto structural connections [1]. An emerging view suggests that structural connection patterns are indeed major constraints for the dynamics of brain activity [13], which are partly captured by functional and effective connectivity. However, whether BOLD propagation is generated only through temporally ordered processes of neuronal origin unfolding through underlying neuroanatomical networks, or also through additional changes in physiological, metabolic or vascular variables remains an issue of debate [54]. Indeed, the precise neuronal mechanisms that determine the spatial and temporal distribution of BOLD signal co-fluctuations and propagation are not yet fully understood. If BOLD fluctuations reflect the consequences of spiking activity, aggregate sub-threshold fluctuations [50], or metabolic relationships among neurons, astrocytes and the supporting capillary network (i.e. neuro-vascular coupling) [62, 63], is an area of ongoing research. On the one hand, BOLD activity propagation patterns within areas during VFM, together with transient, retinotopically selective, co-activation between different visual areas, appears to reflect retinotopically organized switching of spiking input activity between voxels sharing similar visual field position selectivity and tuning characteristics [40, 6, 46, 84]. On the other hand, during RS, BOLD propagation patterns may reflect the footprint of slow subthreshold fluctuations in local field potentials, which can be retinotopically organized and are known to be good predictors of the BOLD signal spatiotemporal covariance structure [51, 12]. Indeed, a recent study by Matsui and colleagues [53] using neuronal calcium signals and simultaneous hemodynamic recordings brings together these lines of evidence by demonstrating that both global fluctuations, in the form of waves propagating across cortex, and transient local co-activations in calcium signals are necessary for setting the spatiotemporal covariance structure of hemodynamic signals [53]. Another line of evidence points to neuronal mechanisms of inter-areal coupling and modulation reflected in the estimated changes in EC. A recent study analyzed simultaneous recordings from V1 and V4 in monkeys and showed that feedforward interactions from V1 to V4 were based on frequencies around the gamma band [42, 56] whereas feedback interactions from V4 to V1 were supported by alpha activity [9, 89]. These study highlights the important fact that different temporal processes may be used as channels over which ‘information’ flows between visual cortical areas. Therefore, care should be taken when interpreting the nature of the estimated interactions in EC.

Furthermore, non-neuronal mechanisms such as the wave-like propagation of hemodynamic activity originating in pial arterioles has been described [70] and related to oscillations in systemic blood pressure (the so called ‘Mayer waves’) [38]. Similarly, large veins draining to the dural sinuses near the occipital pole, which are known to modulate the phase of nearby hemodynamic fluctuations with little effect on signal amplitude [55, 88], may also play a role in shaping, for instance, the periphery-to-fovea interactions observed during RS. Moreover, acting as a temporal low pass filter, neurovascular coupling mechanisms involving the activity of astrocytes [63] and the diffusion of vasodilatory signalling molecules (i.e. nitric oxide) may play an important role in setting the pace of BOLD activity propagation patterns. Together, all these lines of evidence point to the importance of considering the multiple physiological factors implicated in shaping the BOLD signal when interpreting patterns of BOLD activity propagation.

The different propagation patterns of BOLD activity during RS and VFM, as assessed with the present EC analysis, demonstrate distinct cortical dynamics during visual stimulation and in its absence [41, 47]. Nevertheless, in a previous analysis of the present dataset we found that BOLD activity across cortical locations of V1, V2 and V3 sharing similar visual field selectivity (functionally homotopic) can co-fluctuate during RS, enabling the estimation of connective field models from RS data (see figure Figure 3 in Gravel et al. (2014)) [23] that resemble those obtained from VFM data. The fact that retinotopically congruent co-fluctuations in BOLD activity across visual cortical areas can also occur during RS [34, 23, 10] suggests that non-retinal and ‘top-down’ influences, such as feedback modulation of V1 responses, may play a role generating structured patterns of BOLD propagation [59]. A variety of behavioral processes, such as memory consolidation and learning, may recruit V1 into a processing stream, even without external visual stimulation [43, 77, 67]. Together, these studies suggest that, during RS, periods of highly organized neuronal activity in the visual cortex give rise to transitory periods of retinotopically organized BOLD activity propagation. Co-fluctuations within and between early cortical visual field maps may follow, likely reflecting different states of cortical processing [19].

Finally, the current study assesses BOLD propagation in four healthy participants. Although our results are consistent across participants, further studies involving more participants are advised. Moreover, the EC models were estimated based on entire RS and VFM scans. As such, they estimate average BOLD propagation patterns and do not capture specific intervals of variation in these. To establish the neuronal mechanisms underlying the observed changes in EC, further research is still necessary.

## 5 Concluding remarks

We have shown that the propagation of BOLD activity through early visual cortex reveals different directed interaction regimes across the topography and hierarchy of visual cortical areas V1, V2 and V3 during both RS and VFM. We relate these differences in the estimated EC to a task-dependent reconfiguration of feedfoward and feedback interactions throughout the visual system, and changes in *Σ* to a task-dependent neuronal modulation of local cortical excitability. Our results add to a growing body of evidence suggesting that recurrent connectivity and cortico-cortical feedback plays an important role in visual processing. They are consistent with the hypothesis that directed influences (i.e. feedback to V1), as well as intrinsic connectivity (i.e. cortical excitability), interact differently during visual stimulation and rest as a consequence of the visual system using an efficient predictive strategy to process incoming stimuli. We conclude by answering our original question of how the propagation of BOLD activity can also reveal relevant aspects of brain activity at a more local scale. By acknowledging the existence of propagated disturbances in BOLD activity, our approach provides a simple method to infer the local excitability of visual cortical areas and the directed influences unfolding among them during distinct behavioral states.

## 6 Acknowledgments

Acknowledgements Nicolas Gravel was supported by the (Chilean) National Commission for Scientific and Technological Research (BECAS CHILE) and the Graduate School for Medical Sciences (GSMS) of the University Medical Center Groningen (UMCG). Ben M. Harvey, Remco J. Renken and Frans W. Cornelissen were supported by the Netherlands Organization for Scientific Research (NWO Brain and Cognition grant 433-09-233). Ben M. Harvey was also supported by the Portuguese Foundation for Science and Technology (Investigator grant #IF/01405/2014) and by the Netherlands Organization for Scientific Research (VIDI grant 452-17-012).

## 7 Supplementary materials

Video 1S: https://www.dropbox.com/s/qibu1s67e9i7tn3/Video1S.avi?dl=0 Movie illustrating the propagation of BOLD activity during rest depicted in the flattened cortical surface reconstruction of the occipital pole of one participant (total duration: 6 minutes. speed: ×3).

**Fig. S1:**
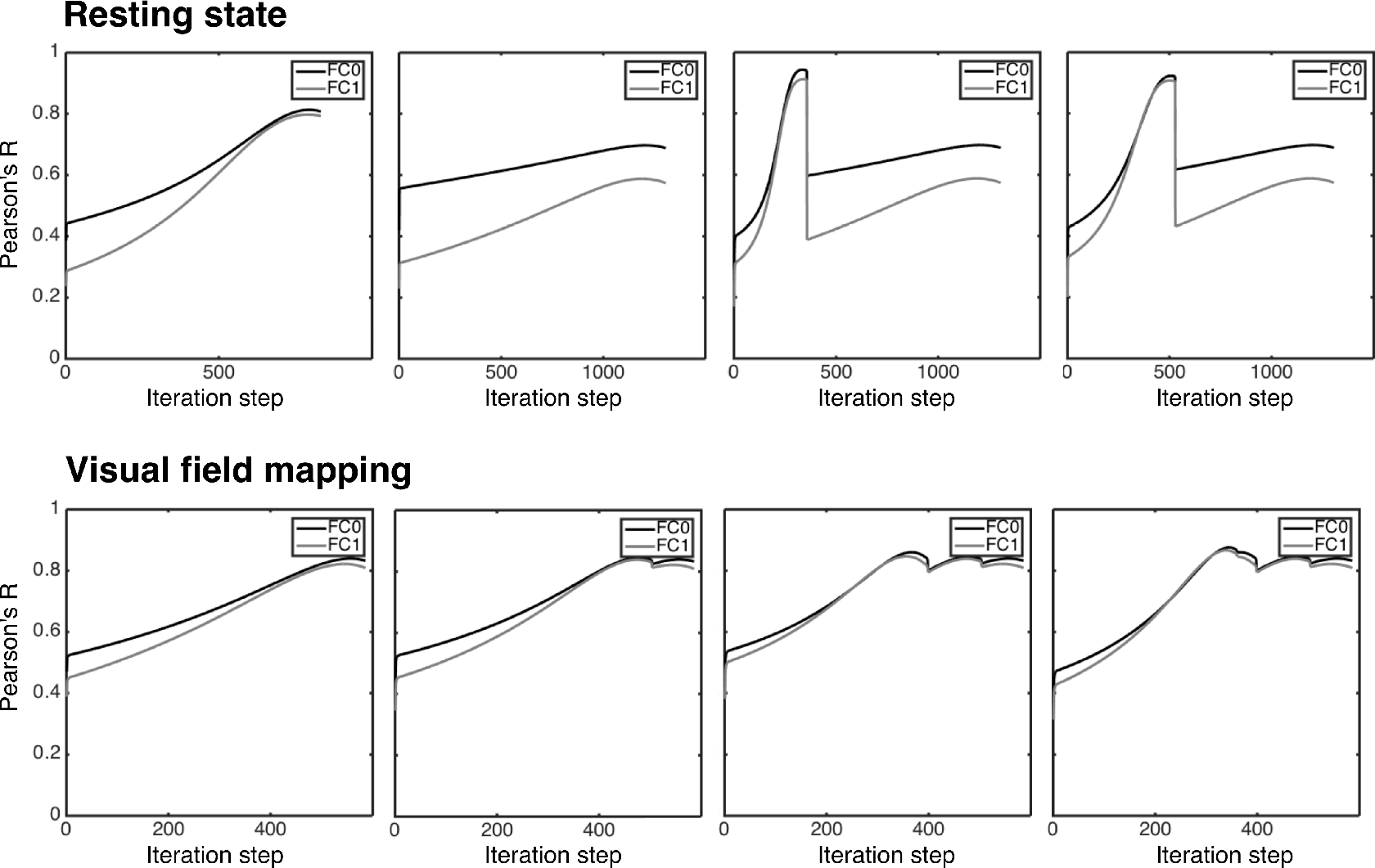
Model fit as a function of iteration step for all participants and conditions. Lines depict the Pearson correlation between the model and the empirical spatiotemporal covariances (FC0 and FC1 in the legend) as a function of the the iteration steps in the Lyapunov optimization.

